# A stress-dependent TRIM28–ALKBH2 feedback loop modulates chemoresistance in NSCLC

**DOI:** 10.1101/2025.11.21.689678

**Authors:** Zhiming Sun, Yunfang Deng, Yue Liu, Zhaohui Liu, Jiabing Li, Lihui Wu, Liyuan Zeng, Xiaorong Feng, Lin Miao, Ying Sheng, Bei Chen, Yuming He, Ye Liu, Yu Zhao

**Affiliations:** The National & Local Joint Engineering Laboratory of Animal Peptide Drug Development, College of Life Sciences, Hunan Normal University, Changsha, Hunan, 410081, China; Peptide and Small Molecule Drug R&D Platform, Furong Laboratory, Hunan Normal University, Changsha, Hunan, 410081, China

**Author notes:** Corresponding author: Yu Zhao.

**Keywords:** TRIM28, ALKBH2, MAGEA6, Ubiquitination, Chemoresistance

## Abstract

Chemoresistance to DNA-damaging agents, including platinum and alkylating compounds, limits treatment efficacy in non-small cell lung cancer (NSCLC) and is frequently associated with elevated DNA repair activity. Here, we identify a stress-responsive feedback loop between the E3 ligase TRIM28 and the demethylase ALKBH2. TRIM28 binds ALKBH2 and promotes its K48-linked polyubiquitination and proteasomal degradation, whereas ALKBH2 enhances TRIM28 transcription and protein levels, forming a feedback loop. Notably, alkylation stress induces a biphasic response: acute MMS exposure enhances TRIM28–ALKBH2 association and accelerates ALKBH2 degradation, whereas prolonged exposure promotes TRIM28 degradation, leading to ALKBH2 stabilization and transcriptional upregulation. Clinically, ALKBH2 is frequently elevated in lung adenocarcinoma and is associated with worse survival, whereas TRIM28 exhibits prognostic value specifically in chemotherapy-treated patients. Functionally, MMS–cisplatin co-treatment increased DNA damage and reduced clonogenic survival by counteracting ALKBH2-dependent alkylation tolerance, linking the TRIM28–ALKBH2 loop to chemoresistance in NSCLC.

## Introduction

Cells are continuously exposed to endogenous metabolites and exogenous alkylating agents that induce cytotoxic and mutagenic DNA adducts, posing a persistent threat to genome integrity (*1, 2*). These lesions are repaired through three principal mechanisms: base excision repair (BER), direct reversal by *O*^6^-methylguanine-DNA methyltransferase (MGMT), and oxidative demethylation catalyzed by AlkB family of Fe(II)/α-ketoglutarate–dependent dioxygenases (*3–5*). AlkB enzymes remove N-linked methyl lesions such as 1-meA and 3-meC via a coupled oxidative reaction that restores the native base (*6–8*). In mammalian cells, these dioxygenases function alongside BER and MGMT to resolve lesions that escape other repair pathways, thereby contributing to genome maintenance across diverse cellular contexts. Changes in AlkB expression or stability are associated with distinct cellular responses to alkylation damage, emphasizing the need for proper regulation of this repair axis.

Although AlkB family enzymes are subject to transcriptional and epigenetic regulation (*9–12*), accumulating evidence indicates that AlkB family dioxygenase protein levels are controlled by ubiquitin-dependent proteostasis. Ubiquitin-regulatory networks have been well characterized for several AlkB family homologues, most notably FTO and ALKBH5. FTO turnover is regulated by multiple context-dependent E3 ligases, including TRIM17, DTX2, MIB1, and RNF144A, and is further modulated by TRIM21 following O-GlcNAc–primed modification (*13–17*). ALKBH5 stability is similarly regulated by the E3 ligase RNF180 and by PRMT5-dependent methylation, the latter promoting its recognition by TRIM28 (*18, 19*). These degradative pathways are counterbalanced by substrate-specific deubiquitinases (DUBs), including USP7 and USP18 for FTO (*20, 21*), and USP14 and USP36 for ALKBH5 (*22, 23*). OTUD4 has also been reported to stabilize the related demethylase ALKBH3 through a noncanonical scaffolding DUB mechanism (*2*), consistent with ubiquitin-dependent regulation observed across multiple AlkB-family enzymes. However, whether this ubiquitin-dependent regulatory paradigm extends to all AlkB family members remains incompletely understood.

Among the nine mammalian AlkB homologues, ALKBH2 functions as a major demethylase involved in the repair of methylated bases in duplex DNA and in limiting alkylation-induced mutagenesis (*24–27*). ALKBH2 is frequently upregulated in cancer, particularly in non–small cell lung cancer (NSCLC), where its elevated expression is associated with increased tolerance to DNA damage and reduced sensitivity to DNA-damaging chemotherapeutic agents, including alkylating and platinum-based drugs (*26, 28–30*). Given that alkylating chemotherapeutics, such as temozolomide, and environmental exposures, including cigarette smoke, generate overlapping classes of methylated DNA lesions (*1, 31, 32*), sustained alkylation stress imposes selective pressure on pathways regulating ALKBH2 protein levels (*10, 33*). However, the ubiquitin-dependent mechanisms regulating ALKBH2 stability—including whether ALKBH2 undergoes ubiquitination and the identity of the E3 ligases or DUBs involved—remain incompletely defined. The ALKBH2-associated ASCC complex recruits TRIM28 upon MMS-induced alkylation stress (*34*), positioning TRIM28 within the alkylation repair context and suggesting a potential regulatory link between TRIM28 and ALKBH2.

The tripartite motif (TRIM) family of RING-type E3 ligases, including TRIM28 and TRIM24, plays a central role in ubiquitin signaling, integrating chromatin regulation with cellular stress responses (*35, 36*). TRIM28 utilizes its N-terminal RING domain and C-terminal PHD–BRD module to mediate KRAB-ZNF–dependent transcriptional silencing, contributing to the maintenance of genomic stability (*37, 38*). This modular architecture enables TRIM28 to function both as a transcriptional corepressor and an E3 ligase, with its activity and chromatin engagement dynamically regulated. In response to DNA double-strand breaks, TRIM28 undergoes ATM/ATR-dependent phosphorylation at Ser824, which relaxes heterochromatin and facilitates the recruitment of repair factors (*39, 40*). TRIM28 expression is further regulated by transcriptional and post-transcriptional programs, as well as by ubiquitin-dependent turnover under genotoxic or metabolic stress (*41–43*). Following DNA double-strand breaks, RNF4 targets TRIM28 for SUMOylation-dependent degradation (*44*). In parallel, under metabolic stress, March5 promotes TRIM28 ubiquitination and reduces its protein stability (*45*). Moreover, MAGE family proteins, including MAGEA6, serve as cofactors that promote TRIM28 recruitment to its substrates by facilitating the assembly of TRIM28–substrate E3 complexes (*46, 47*). Although multiple regulatory layers of TRIM28 have been described, whether and how TRIM28 participates in alkylation damage repair, and how its regulation is modulated under alkylating stress, remain poorly understood.

In this study, we identify a stress-responsive feedback loop involving TRIM28, its cofactor MAGEA6, and the DNA demethylase ALKBH2. TRIM28 binds ALKBH2 and promotes its K48-linked polyubiquitination and proteasomal degradation, whereas ALKBH2 reciprocally enhances TRIM28 transcription and protein levels. This loop is modulated by alkylation damage in a time-dependent manner: short-term alkylation exposure enhances TRIM28–ALKBH2 association and accelerates ALKBH2 degradation, whereas prolonged MMS exposure promotes TRIM28 ubiquitination and degradation, leading to ALKBH2 stabilization and enhanced transcription. These data offer insights into how alkylation stress duration and context regulate ALKBH2 availability and modulate DNA repair capacity and chemoresistance in NSCLC.

## Results

### TRIM28 directly binds ALKBH2

To explore proteins associated with TRIM28, we performed tandem affinity purification in H1299 non–small cell lung cancer (NSCLC) cells, followed by silver staining and LC–MS/MS analysis. ALKBH2 was identified as a candidate TRIM28-associated protein (Fig. 1A, fig. S1A, and table S1). Co-immunoprecipitation (co-IP) assays in HEK293T cells coexpressing HA-tagged ALKBH2 and Flag-tagged TRIM28 confirmed this interaction, whereas no detectable association was observed with the related paralog TRIM24 (Fig. 1B). Reciprocal co-IP using HA-ALKBH2 further validated the specificity of the interaction (Fig. 1C).

**Figure 1.**
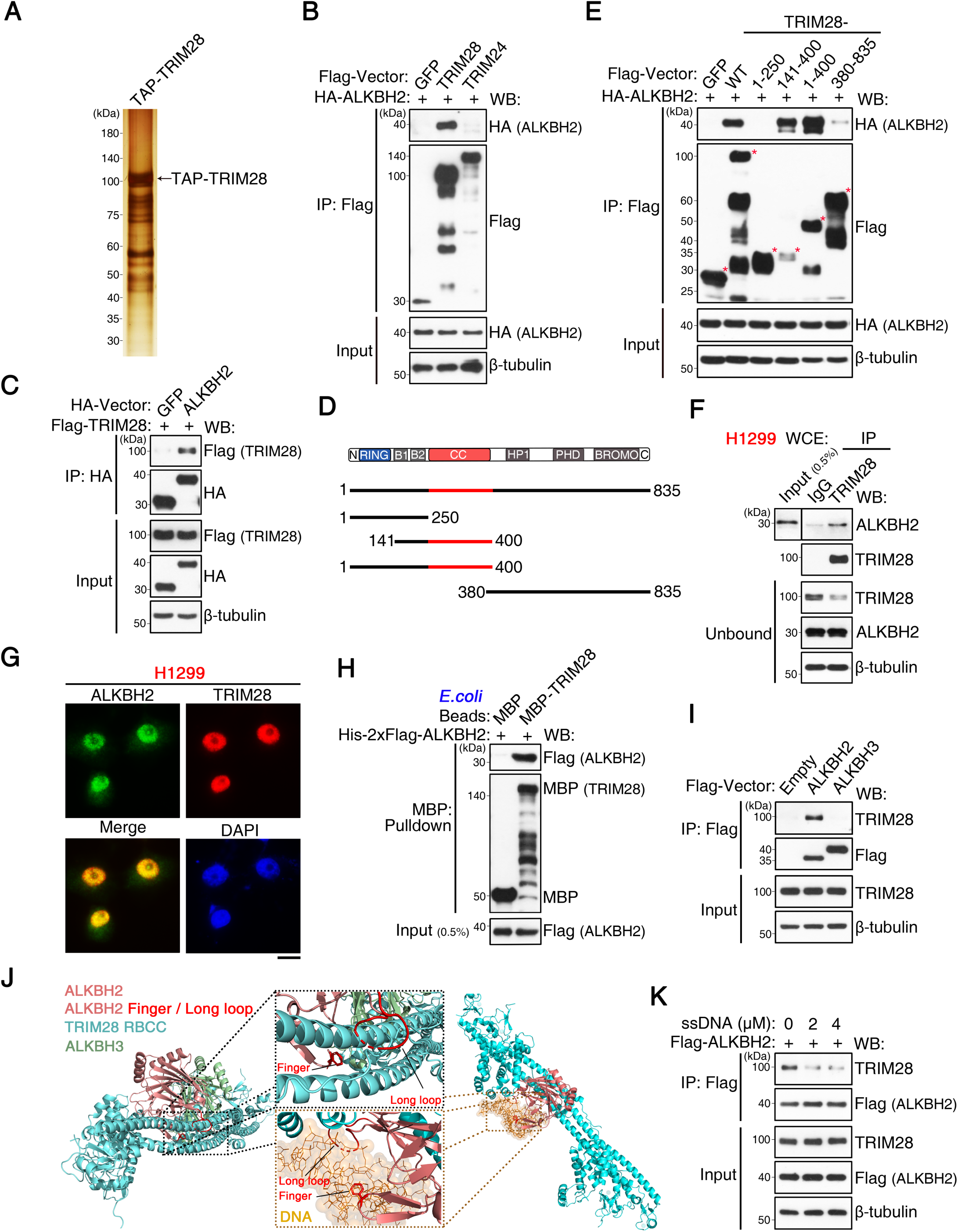
TRIM28 directly binds ALKBH2. **(A)** Silver staining and LC–MS/MS identification of ALKBH2 as a TRIM28-associated protein following affinity purification from H1299 NSCLC cells; the TRIM28 band is indicated. **(B)** Co-immunoprecipitation (co-IP) analysis of HEK293T cells co-expressing HA-ALKBH2 and Flag-TRIM28 or Flag-TRIM24; Flag IP was followed by western blotting (WB). Data are representative of two independent experiments with similar results. **(C)** Reciprocal co-IP using HA-ALKBH2 and Flag-TRIM28 (or HA-GFP control), HA IP was followed by WB. Data are representative of two independent experiments with similar results. **(D)** Schematic of full-length TRIM28 and truncation mutants. **(E)** Domain mapping using Flag IP followed by WB in HEK293T cells co-transfected with HA-ALKBH2 and individual TRIM28 truncation constructs. Data are representative of two independent experiments. **(F)** Endogenous ALKBH2–TRIM28 interaction detected by co-IP using an anti-TRIM28 antibody in H1299 cell lysates. Data are representative of two independent experiments. **(G)** Immunofluorescence (IF) showing nuclear co-localization of endogenous ALKBH2 (green) and TRIM28 (red); nuclei were counterstained with DAPI (blue). Scale bar, 20 µm. Pearson correlation coefficient (PCC) = 0.896 ± 0.016 (mean ± SD, n = 42 cells). **(H)** *In vitro* MBP pull-down assay using recombinant His-Flag-ALKBH2 and purified MBP-TRIM28. Data are representative of two independent experiments. **(I)** Specificity profiling of endogenous TRIM28 binding in HEK293T cells transfected with Flag-ALKBH2 or Flag-ALKBH3. Data are representative of two independent experiments with similar results. **(J)** GRAMM-X docking of TRIM28 RBCC (cyan) with ALKBH2 (light red) or ALKBH3 (green). Middle panel: magnified interface highlighting the Finger and Long-loop motifs (red). Right panel: superposition with the ALKBH2–DNA structure (orange), showing partial overlap between TRIM28- and DNA-binding surfaces. **(K)** Flag IP of lysates from HEK293T cells expressing Flag-ALKBH2 in the presence of increasing ssDNA concentrations. Data are representative of two independent experiments with similar results.

To determine the TRIM28 region required for ALKBH2 binding, we performed domain-mapping analyses using a series of TRIM28 truncation mutants. These experiments indicated that the BCC region (B-box plus coiled-coil; residues 141–400) is required for ALKBH2 association (Fig. 1, D and E). ALKBH2 co-precipitated with TRIM28 in endogenous co-IP experiments, indicating that the interaction occurs under physiological conditions (Fig. 1F). Moreover, immunofluorescence analysis revealed prominent nuclear colocalization of TRIM28 and ALKBH2 (Fig. 1G).

*In vitro* MBP pull-down assays using purified recombinant proteins showed that TRIM28 directly binds ALKBH2 (Fig. 1H). Specificity assays revealed stronger binding to ALKBH2 than to ALKBH3 (Fig. 1I) or the direct-reversal repair enzyme MGMT (fig. S1B). Protein–protein docking predicted a TRIM28–ALKBH2 interface that overlaps the nucleic-acid-binding region of ALKBH2, centered on the Finger (hairpin) motif and the flexible Long-loop (*48*), which are present in ALKBH2 but absent in ALKBH3 (Fig. 1J). Consistent with this model, single-stranded DNA (ssDNA) competitively reduced TRIM28–ALKBH2 binding in a dose-dependent manner (Fig. 1K), a commonly used approach to assess the DNA-occupied states of repair enzymes (*49–51*). Collectively, these data indicate that TRIM28 directly binds ALKBH2 through its BCC domain and propose a mechanism in which DNA binding limits TRIM28 access to the ALKBH2 interaction interface.

### A reciprocal TRIM28–ALKBH2 regulatory loop coordinates ALKBH2 ubiquitination and stability, with ALKBH2 enhancing TRIM28 transcription

To determine whether ALKBH2 undergoes ubiquitin-dependent modification, we performed denaturing immunoprecipitation (IP) of Flag-ALKBH2 from HEK293T cells co-transfected with HA-ubiquitin (Ub). A high-molecular-weight smear corresponding to polyubiquitinated ALKBH2 was detected and was markedly increased upon proteasome inhibition with MG132 (Fig. 2A). Expression of a K48-only ubiquitin mutant further enhanced ALKBH2 ubiquitination, whereas the K48R ubiquitin mutant almost completely abolished it (Fig. 2B), suggesting that ALKBH2 is primarily modified by K48-linked polyubiquitin chains. Halo-TUBE pull-down assays confirmed that endogenous ALKBH2 undergoes K48-linked polyubiquitination (Fig. 2C).

**Figure 2.**
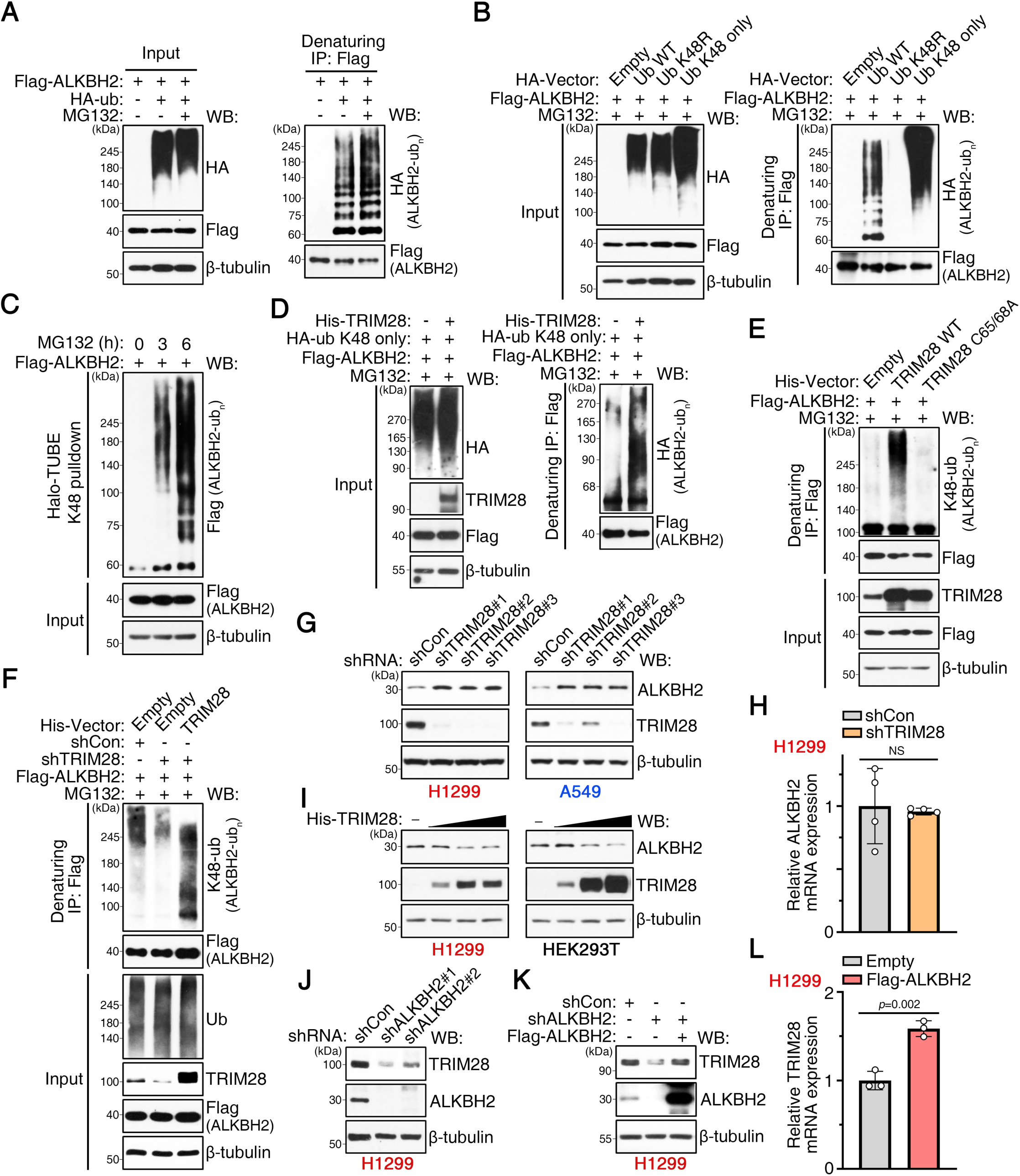
TRIM28 promotes K48-linked ubiquitination and proteasomal degradation of ALKBH2. **(A)** In vivo ubiquitination assay in HEK293T cells co-transfected with Flag-ALKBH2 and HA-ubiquitin (Ub), treated with MG132 (10 µM, 6 h). Lysates were subjected to denaturing Flag IP (1% SDS) followed by WB. Data are from three independent experiments. **(B)** Denaturing Flag IP followed by WB in HEK293T cells co-transfected with Flag-ALKBH2 and HA-tagged WT, K48-only, or K48R ubiquitin. Data are representative of two independent experiments with similar results. **(C)** K48-linked polyubiquitin enrichment using Halo-TUBE beads in HEK293T cells expressing Flag-ALKBH2 treated with MG132. Data are representative of two independent experiments with similar results. **(D)** Denaturing Flag IP in HEK293T cells co-transfected with Flag-ALKBH2, His-TRIM28, and HA-K48-only ubiquitin. Data are representative of two independent experiments with similar results. **(E)** Analysis of ALKBH2 ubiquitination in HEK293T cells co-transfected with Flag-ALKBH2 and WT or RING-inactive (C65/68A) His-TRIM28. Representative of two independent experiments. **(F)** Denaturing Flag IP assessing K48-linked ALKBH2 ubiquitination in TRIM28-knockdown HEK293T cells reconstituted with shRNA-resistant TRIM28. Representative of two independent experiments. **(G)** WB analysis of H1299 and A549 cells transduced with control (shCon) or TRIM28 shRNAs. N = 3 independent experiments. **(H)** qRT-PCR quantification of ALKBH2 mRNA in H1299 cells after TRIM28 depletion. Data points are shown as technical replicates (N = 4); experiments were independently repeated twice with similar results. **(I)** WB analysis of ALKBH2 levels in H1299 and HEK293T cells overexpressing graded doses of TRIM28. N = 3 independent repeats. **(J)** WB analysis of H1299 cells transduced with control or ALKBH2 shRNAs. N = 3 independent experiments. **(K)** WB analysis of ALKBH2-depleted H1299 cells reconstituted with Flag-ALKBH2. N = 3 independent repeats. **(L)** qRT-PCR of TRIM28 transcript levels in H1299 cells expressing empty vector or Flag-ALKBH2 (N = 4). Data were obtained using the same replication scheme as in (H). Error bars indicate mean ± SD. Statistical comparisons were performed using unpaired two-tailed Student’s *t*-tests; NS, not significant.

We next examined whether TRIM28 mediates ALKBH2 ubiquitination. Overexpression of wild-type (WT) TRIM28 increased K48-linked ubiquitination of ALKBH2 (Fig. 2D), whereas the RING-inactive C65/68A mutant, which abolishes TRIM28 E3 ligase activity (*46*), did not enhance ALKBH2 ubiquitination (Fig. 2E). TRIM28 depletion using two independent shRNAs markedly reduced ALKBH2 ubiquitination (fig. S2A), and re-expression of shRNA-resistant TRIM28 restored this activity (Fig. 2F). These data indicate that TRIM28 E3 ligase activity is required to promote efficient K48-linked ubiquitination of ALKBH2.

We then examined whether TRIM28 regulates ALKBH2 protein stability. TRIM28 knockdown in H1299 and A549 cells increased ALKBH2 protein levels without altering its mRNA expression (Fig. 2, G and H). This effect was reproduced by CRISPR/Cas9-mediated TRIM28 depletion (fig. S2B). Conversely, increasing TRIM28 expression led to a dose-dependent decrease in ALKBH2 protein levels in both H1299 and HEK293T cells (Fig. 2I). Cycloheximide (CHX) chase assays showed that TRIM28 knockout extended the half-life of ALKBH2, whereas TRIM28 overexpression shortened it in a RING-dependent manner (fig. S2, C to F). Notably, these changes occurred without altering ALKBH2 subcellular localization (fig. S2G). These data indicate that TRIM28 regulates ALKBH2 protein stability by promoting its degradation.

To determine whether ALKBH2 in turn regulates its E3 ligase, we depleted ALKBH2 in H1299 cells using two independent shRNAs. TRIM28 protein levels were reduced following ALKBH2 knockdown (Fig. 2J), and re-expression of Flag-ALKBH2 restored TRIM28 protein levels (Fig. 2K). In parallel, ALKBH2 overexpression increased TRIM28 mRNA levels as measured by qRT-PCR (Fig. 2L). Consistent with this regulatory relationship, analysis of TCGA lung adenocarcinoma datasets revealed a positive correlation between ALKBH2 and TRIM28 transcript levels (fig. S2H). Collectively, these data identify a feedback loop in which TRIM28 promotes K48-linked ubiquitination and proteasome-dependent degradation of ALKBH2, whereas ALKBH2 enhances TRIM28 transcription and protein levels.

### MAGEA6 enhances TRIM28 E3 ligase assembly and activity toward ALKBH2, a function impaired by cancer-associated variants

MAGE family proteins interact with TRIM-type E3 ligases to modulate their activity (*46, 52*). To assess how MAGEA6 contributes to the organization of the TRIM28–ALKBH2 complex, we examined whether MAGEA6 associates with ALKBH2. Co-IP in HEK293T cells revealed an association between MAGEA6 and ALKBH2 (Fig. 3A), and pull-down assays using purified proteins confirmed a direct interaction (Fig. 3B). TRIM28 depletion did not affect MAGEA6–ALKBH2 binding (Fig. 3C), suggesting that this interaction occurs independently of TRIM28. MAGEA6 overexpression enhanced TRIM28–ALKBH2 association in a dose-dependent manner (Fig. 3D and fig. S3A), whereas knockdown of endogenous MAGEA3/6 attenuated this interaction (Fig. 3E); MAGEA3 and MAGEA6 were analyzed collectively due to their 96% sequence homology (*53*). Furthermore, the related protein MAGEC2 also enhanced TRIM28–ALKBH2 assembly (fig. S3B), consistent with a conserved role of MAGEA/C family members in orchestrating this complex.

**Figure 3.**
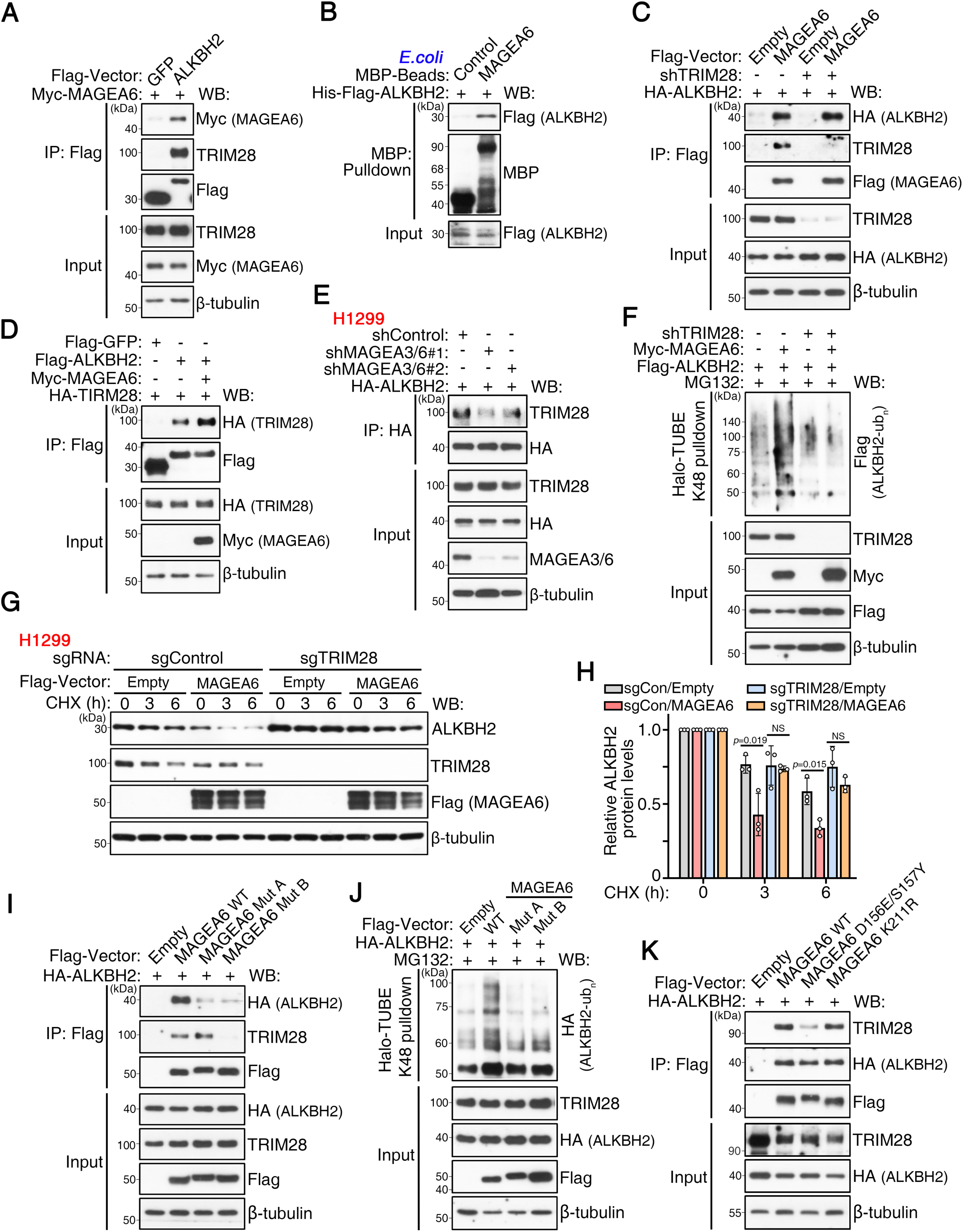
MAGEA6 enhances the TRIM28–ALKBH2 interaction and promotes ALKBH2 degradation. **(A)** Flag IP analysis of the interaction between Myc-MAGEA6 and Flag-ALKBH2 (or GFP control) in HEK293T cells. Representative of two independent experiments with similar results. **(B)** *In vitro* pull-down assay using recombinant His-Flag-ALKBH2 and purified MBP-MAGEA6. Representative of two independent experiments. **(C)** Flag IP in HEK293T cells (± TRIM28 knockdown) co-transfected with HA-ALKBH2 and Flag-MAGEA6. Representative of two independent experiments. **(D)** Flag IP analysis of the ALKBH2–TRIM28 interaction in HEK293T cells transfected with Myc-MAGEA6. Representative of two independent experiments. **(E)** HA IP in H1299 cells stably expressing HA-ALKBH2 following MAGEA3/6 knockdown with shMAGEA3/6#1 or shMAGEA3/6#2. Representative of two independent experiments with similar results. **(F)** Halo-TUBE pull-down in TRIM28-knockdown HEK293T cells overexpressing Myc-MAGEA6. Representative of two independent experiments. **(G)** WB analysis of ALKBH2 stability in sgControl or TRIM28-knockout (sgTRIM28) H1299 cells transfected with MAGEA6 and treated with cycloheximide (CHX, 150 µg/mL). **(H)** Quantification of ALKBH2 protein levels from (G). Data were quantified from 3 biological replicates (N = 3). **(I)** Flag IP mapping interactions between Flag-MAGEA6 (WT or mutants), endogenous TRIM28, and co-expressed HA-ALKBH2 in HEK293T cells. Representative of two independent experiments. **(J)** Halo-TUBE pull-down of ubiquitinated proteins in HEK293T cells co-expressing HA-ALKBH2 and Flag-MAGEA6 (WT or mutants). Representative of two independent experiments with similar results. **(K)** Flag IP of HEK293T cells transfected with HA-ALKBH2 and cancer-associated Flag-MAGEA6 mutants. Representative of two independent experiments. Error bars indicate mean ± SD. Statistical comparisons were performed using unpaired two-tailed Student’s *t*-tests. NS, not significant.

We next examined whether MAGEA6 regulates TRIM28-dependent ubiquitination and turnover of ALKBH2. MAGEA6 knockdown reduced endogenous ALKBH2 ubiquitination, whereas re-expression of MAGEA6 reversed this modification (fig. S3C). Conversely, MAGEA6 overexpression promoted K48-linked ubiquitination of ALKBH2, an effect that was abolished upon TRIM28 depletion (Fig. 3F), indicating a TRIM28-dependent requirement for MAGEA6 in ALKBH2 ubiquitination. Notably, MAGEA6 promoted ALKBH2 degradation in control cells, whereas this effect was not observed in TRIM28-knockout H1299 cells (Fig. 3, G and H).

Cancer-associated variants in MAGEA6 have been reported across multiple tumor types, including lung cancer (*54*). We therefore analyzed endogenous MAGEA6 transcripts in H1299 cells and identified two variants, MutA and MutB, harboring 12 and 20 amino acid substitutions, respectively (Fig. S3D). Both variants were cloned and sequence-verified. Co-IP assays revealed that both MutA and MutB showed reduced interaction with ALKBH2, whereas MutB additionally reduced interaction with TRIM28 (Fig. 3I). Consistent with these binding defects, both variants were less effective than wild-type MAGEA6 at enhancing ALKBH2 ubiquitination and degradation (Fig. 3J and fig. S3E).

To explore the structural basis of this interaction, AlphaFold3 modeling identified a putative TRIM28-binding interface within the MAGE homology domain involving residues D156 and S157 (fig. S3F). Within this interface, S157 is substituted in the MutB variant and overlaps with tumor-associated substitutions reported in COSMIC for MAGEA6 and the homologous MAGEA3. Structure-guided mutagenesis further showed that an interface-disrupting D156E/S157Y double substitution reduced association with TRIM28 and decreased MAGEA6-dependent ALKBH2 stability (Fig. 3K and fig. S3G). In contrast, the K211R substitution—located within the predicted ALKBH2-binding surface—had minimal effect on MAGEA6–ALKBH2 interaction (Fig. 3K). Collectively, these data indicate that residues D156 and S157 contribute to the MAGEA6–TRIM28 interface and that disruption of this interface impairs MAGEA6-dependent ALKBH2 ubiquitination and degradation. This conclusion is consistent with the enrichment of tumor-associated variants at this site.

### MMS drives a biphasic ALKBH2 response, from acute degradation to prolonged accumulation

#### MMS acutely promotes ALKBH2 degradation

To investigate how MMS regulates the TRIM28–ALKBH2 axis, we examined their association dynamics in H1299 cells. Short-term MMS exposure (3–6 h) increased the TRIM28–ALKBH2 association in a time- and dose-dependent manner (Fig. 4, A and B). Both TRIM28 and ALKBH2 were predominantly nuclear regardless of MMS treatment (fig. S4A). In contrast, MAGEA3/6 showed increased nuclear localization upon MMS exposure (Fig. 4C), consistent with the involvement of MAGEA3/6 in enhanced TRIM28–ALKBH2 assembly during genotoxic stress. Accordingly, MMS increased ALKBH2 K48-linked ubiquitination in a concentration-dependent manner (Fig. 4D).

**Figure 4.**
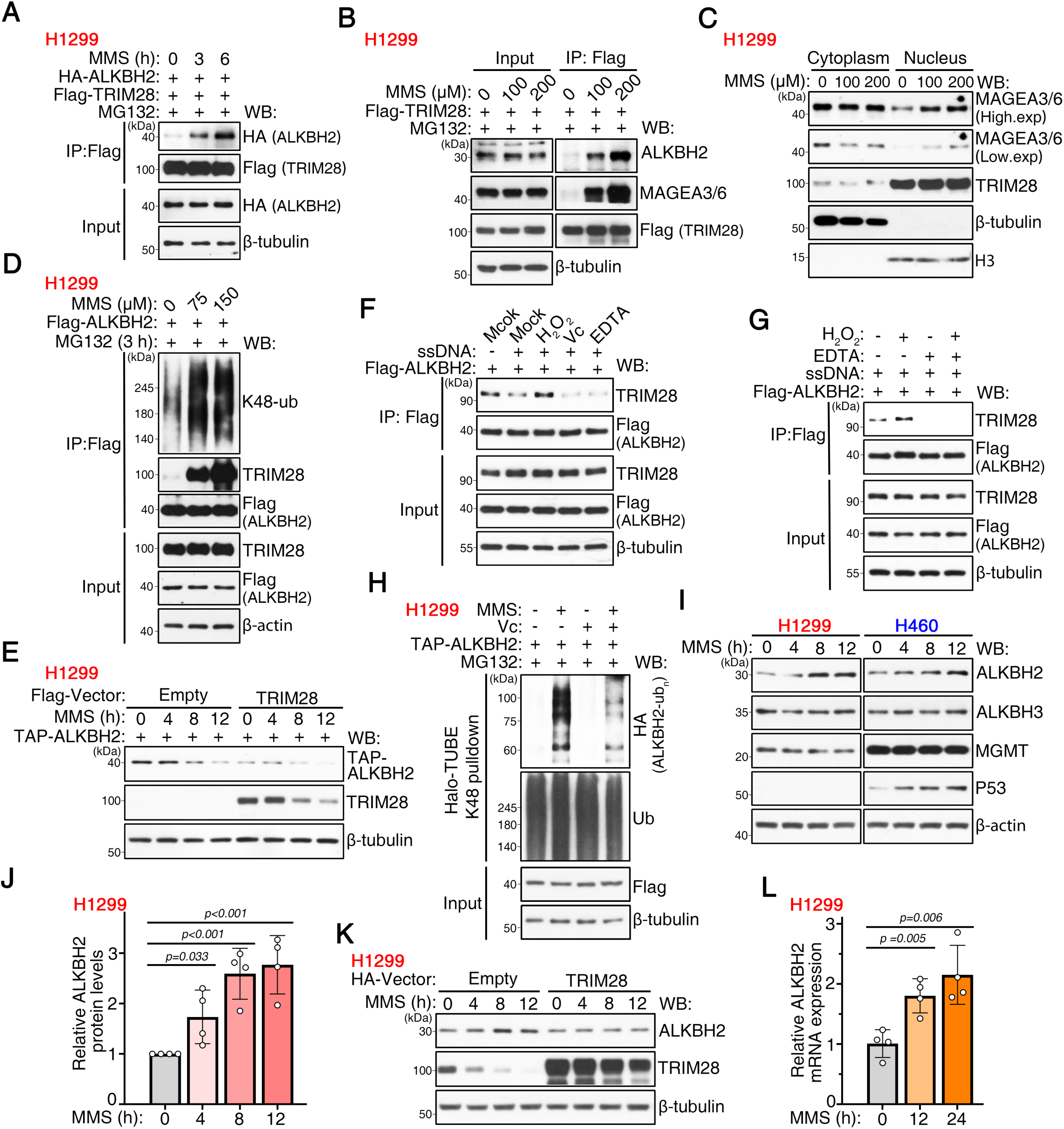
MMS dynamically regulates the TRIM28–ALKBH2 pathway. **(A)** Flag IP from H1299 cells expressing HA-ALKBH2 and Flag-TRIM28 treated with MMS (100 µM) and MG132 (10 µM) for the indicated times. Representative of two independent experiments with similar results. **(B)** Flag IP from H1299 cells expressing Flag-TRIM28 treated with the indicated MMS concentrations (3 h, +MG132). Representative of two independent experiments. **(C)** WB analysis of MAGEA3/6 in nuclear and cytoplasmic fractions of H1299 cells treated with MMS (100 µM, 4 h). Representative of two independent experiments. **(D)** Flag IP of H1299 cells expressing Flag-ALKBH2 treated with MMS (0–200 µM, 3 h) plus MG132. Representative of two independent experiments with similar results. **(E)** Degradation assay in H1299 cells expressing TAP-ALKBH2 transduced with Flag-TRIM28 or vector and treated with MMS (100 µM). N = 3 independent repeats. **(F)** Flag IP of cell lysates incubated with ssDNA (1 µM), H₂O₂ (200 µM), ascorbate (Vc, 500 µM), or EDTA (5 mM). Representative of two independent experiments with similar results. **(G)** Flag IP assessing the effect of EDTA (5 mM) on H₂O₂-induced interactions in ssDNA-containing lysates. Representative of two independent experiments. **(H)** Halo-TUBE pull-down in H1299 cells expressing TAP-ALKBH2 treated with MG132 (10 µM), MMS (100 µM), and Vc (500 µM) for 8 h. Representative of two independent experiments with similar results. **(I)** WB analysis of ALKBH2 in H1299 and H460 cells treated with MMS (100 µM). **(J)** Quantification of ALKBH2 protein levels from (I). Data were quantified from 4 biological replicates (N = 4). **(K)** WB analysis of H1299 cells expressing HA-TRIM28 treated with MMS (100 µM). N = 3 independent experiments. **(L)** qRT-PCR of ALKBH2 in H1299 cells treated with MMS (100 µM). Data points are shown as technical replicates (N = 4); experiments were independently repeated twice with similar results. Error bars indicate mean ± SD. Statistical comparisons were performed using unpaired two-tailed Student’s *t*-tests.

To exclude transcriptional effects, we used H1299 cells stably expressing TAP-tagged ALKBH2 (Flag–HA tandem affinity tag). MMS accelerated ALKBH2 degradation, and this effect was further potentiated by TRIM28 overexpression (Fig. 4E). Endogenous ALKBH2 similarly showed reduced stability when cells were co-treated with MMS and cycloheximide (CHX), compared with CHX alone (fig. S4, B and C). These data indicate an early response phase in which MMS triggers TRIM28-dependent ubiquitination and degradation of ALKBH2.

Because alkylating agents such as MMS and temozolomide (TMZ) elevate intracellular ROS (*55, 56*), we next examined whether redox conditions influence TRIM28–ALKBH2 association. Co-IP analysis revealed that H₂O₂ enhanced this association, whereas ssDNA, ascorbate (Vc), or EDTA reduced it (Fig. 4F). Reduced glutathione (GSH) diminished TRIM28–ALKBH2 association, whereas oxidized glutathione (GSSG) had minimal effect; GSH also attenuated the H₂O₂-induced increase (fig. S4D). EDTA similarly lowered basal association and blocked H₂O₂-driven enhancement (Fig. 4G). Notably, oxidative stress alone was sufficient to induce K48-linked polyubiquitination of ALKBH2 (fig. S4E).

The catalytically inactive ALKBH2(H171A/D173A) mutant (*48, 57*), within the Fe(II)-binding active site, was refractory to both the ssDNA-dependent reduction and the H₂O₂-induced enhancement of TRIM28 binding, indicating that these redox-associated effects depend on ALKBH2 catalytic activity (fig. S4F). The reducing agent Vc further suppressed MMS-induced K48-linked ubiquitination of ALKBH2 (Fig. 4H). Together, these data indicate an acute, redox-responsive phase in which TRIM28-mediated ubiquitination drives ALKBH2 degradation.

### Prolonged MMS exposure stabilizes ALKBH2

In contrast to the early degradative phase, prolonged MMS exposure (≥8 h) progressively increased ALKBH2 protein levels in H1299 and H460 cells (Fig. 4, I and J). In comparison, the related demethylase ALKBH3 and MGMT showed no detectable change in protein levels under the same conditions. TRIM28 overexpression attenuated the MMS-induced increase in ALKBH2 (Fig. 4K). Together, these results indicate that prolonged MMS exposure stabilizes ALKBH2.

### Prolonged MMS exposure induces transcriptional upregulation of ALKBH2

To determine whether ALKBH2 accumulation involves transcriptional activation, we measured ALKBH2 transcript levels in H1299 cells after MMS exposure for the indicated durations. ALKBH2 mRNA increased at 12 h and remained elevated at 24 h (Fig. 4L), paralleling the delayed increase in ALKBH2 protein levels.

Collectively, these data reveal a stress-dependent feedback loop, with an early TRIM28-dependent ubiquitination and degradation phase, followed by a later phase characterized by ALKBH2 stabilization accompanied by transcriptional induction.

### MMS induces K48-linked ubiquitination and proteasomal degradation of TRIM28, coordinated by TRIM24 and MAGEA6

To investigate how TRIM28 is regulated during alkylation stress, we examined its stability and post-translational modification. MMS treatment caused a time- and dose-dependent decrease in TRIM28 protein without altering its mRNA (Fig. 5, A to C), and this reduction was blocked by MG132, consistent with proteasome-mediated degradation. Prolonged low-dose MMS exposure (100 µM, 24–72 h) led to a progressive increase in ALKBH2 protein levels, whereas ALKBH3, MGMT, and MAGEA3/6 remained stable (fig. S5A). A 12-h MMS pre-exposure caused TRIM28 depletion, increased basal ALKBH2 levels, and slower degradation of ALKBH2 (fig. S5, B and C), indicating that reduced TRIM28 availability contributes to ALKBH2 accumulation during sustained alkylation stress.

**Figure 5.**
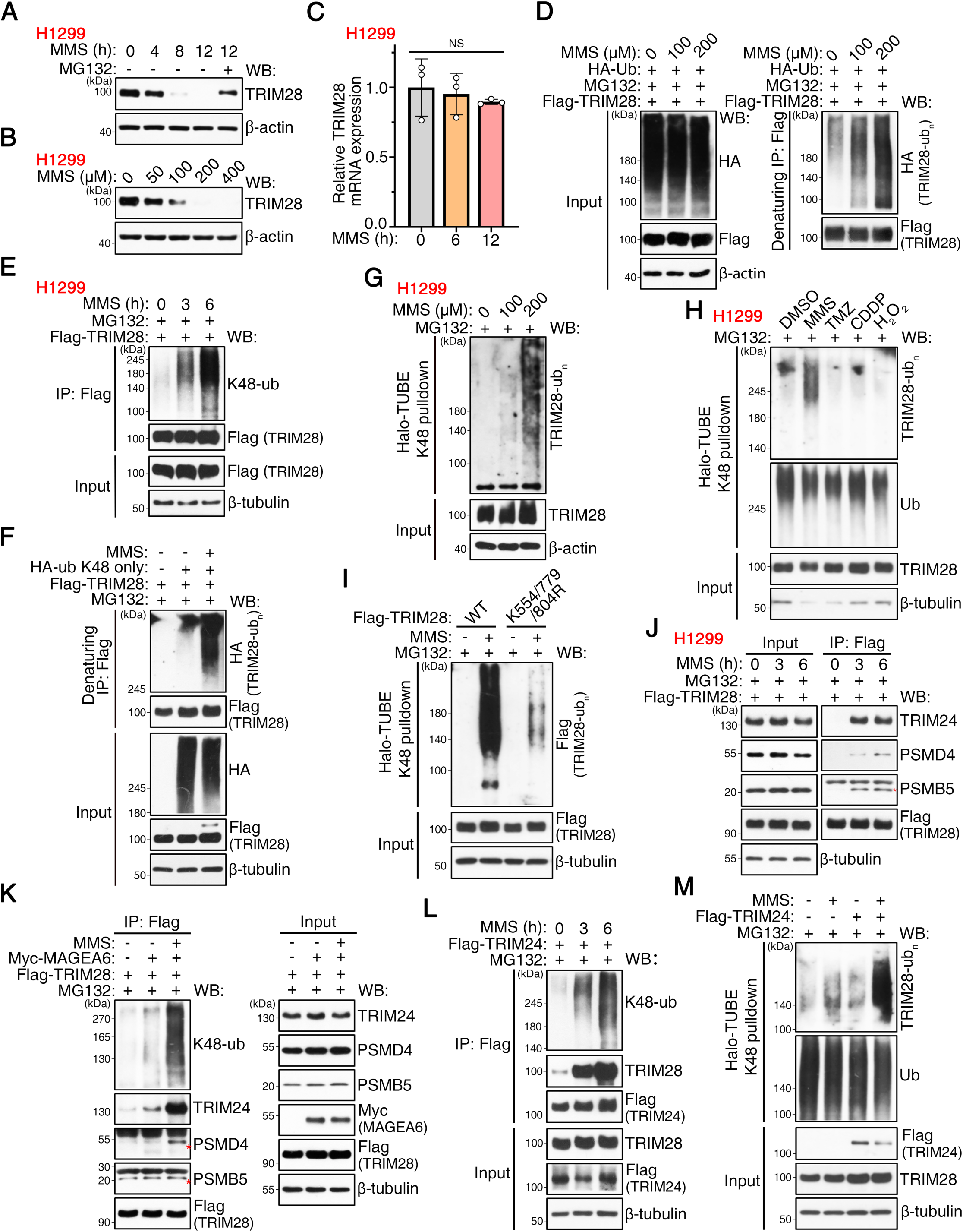
MMS induces K48-linked ubiquitination and proteasomal degradation of TRIM28. (**A, B)** WB analysis of TRIM28 levels in H1299 cells treated with MMS in a time-dependent (A) or dose-dependent (B) manner. N = 3 independent experiments. **(C)** qRT-PCR of TRIM28 mRNA in cells treated with MMS (100 µM). Data points are shown as technical replicates (N = 3); experiments were independently repeated twice with similar results. **(D)** Denaturing Flag IP of HEK293T cells expressing Flag-TRIM28 and HA-Ub treated with MMS and MG132 (10 µM). Representative of two independent experiments with similar results. **(E)** Flag IP and K48-specific ubiquitin WB in H1299 cells expressing Flag-TRIM28 treated with MMS (100 µM) + MG132. Representative of two independent experiments with similar results. **(F)** Denaturing Flag IP in HEK293T cells transfected with Flag-TRIM28 and HA-Ub (K48-only) treated with MMS (100 µM). Representative of two independent experiments. **(G)** Halo-TUBE pull-down of endogenous TRIM28 in H1299 cells treated with MG132 and increasing MMS. Representative of two independent experiments. **(H)** Halo-TUBE pull-down comparing the effects of MMS (200 µM), temozolomide (TMZ, 200 µM), cisplatin (CDDP, 20 µM), or H₂O₂ (300 µM) on TRIM28 ubiquitination (+MG132, 6 h). Representative of two independent experiments with similar results. **(I)** Halo-TUBE pull-down in HEK293T cells expressing Flag-TRIM28 WT or the SUMO-deficient mutant (K554/779/804R) treated with MMS. Representative of two independent experiments. **(J)** Flag IP in H1299 cells expressing Flag-TRIM28 treated with MMS (100 µM) + MG132. Representative of two independent experiments. **(K)** Flag IP in HEK293T cells expressing Flag-TRIM28 and Myc-MAGEA6 treated with MMS. Representative of two independent experiments. **(L)** Flag IP and WB in HEK293T cells expressing Flag-TRIM24 treated with MMS + MG132. Results were reproduced in independent experiments. **(M)** Halo-TUBE enrichment of K48-linked proteins in HEK293T cells overexpressing Flag-TRIM24 treated with MMS. Representative of two independent experiments with similar results. Error bars indicate mean ± SD. Statistical comparisons were performed using unpaired two-tailed Student’s *t*-tests. NS, not significant.

To examine the mechanism underlying this regulation, we analyzed TRIM28 ubiquitination. Denaturing IP of Flag-TRIM28 revealed that MMS markedly induced TRIM28 polyubiquitination, which increased over time in H1299 cells (Fig. 5, D and E). Use of a K48-only ubiquitin mutant indicated that these conjugates are predominantly K48-linked (Fig. 5F). K48-specific Halo-TUBE pulldown detected MMS-induced ubiquitination of endogenous TRIM28 (Fig. 5G). In contrast, other genotoxic agents—including TMZ, CDDP, and H₂O₂—did not induce detectable K48-linked ubiquitination or TRIM28 degradation (Fig. 5H and fig. S5, D to F), indicating that TRIM28 degradation is selective for MMS-induced alkylation stress. The SUMO-acceptor-site TRIM28 mutant (K554/779/804R) (*58*) exhibited impaired K48-linked ubiquitination (Fig. 5I), indicating that this lysine cluster is required for efficient MMS-induced modification.

To identify upstream factors regulating TRIM28 turnover, we analyzed the TRIM28 interactome (table S2) and validated selected interacting proteins. MMS increased TRIM28 association with the E3 ligase TRIM24 and the proteasome subunits PSMD4 and PSMB5 in a time-dependent manner (Fig. 5J), whereas these interactions were weakened in the SUMO-acceptor mutant (fig. S5G). MAGEA6 further enhanced the assembly of the TRIM28–TRIM24 complex. Upon MMS exposure, MAGEA6 co-expression increased TRIM28 binding to TRIM24 and PSMD4/PSMB5, accompanied by enhanced K48-linked TRIM28 ubiquitination (Fig. 5K). Reciprocal co-IP using Flag-TRIM24 confirmed the MMS-inducible TRIM28–TRIM24 interaction and revealed that TRIM24 itself undergoes K48-linked ubiquitination (Fig. 5L). Overexpression of TRIM24 enhanced MMS-induced ubiquitination and accelerated the reduction of endogenous TRIM28 (Fig. 5M and fig. S5, H and I). Collectively, these data indicate that MMS induces K48-linked ubiquitination and proteasomal degradation of TRIM28 via a pathway involving MAGEA6, TRIM24, and SUMO-dependent regulation.

### TRIM28–ALKBH2 axis links patient outcome to alkylation-stress responses

To assess the clinical relevance of this pathway in lung adenocarcinoma (LUAD), we examined expression patterns of key alkylation repair enzymes. Analysis of TCGA transcriptomic and CPTAC proteomic datasets indicated increased ALKBH2 and decreased ALKBH3 in tumors relative to normal lung tissue, whereas MGMT showed no significant change (Fig. 6A and fig. S6A). Immunohistochemistry further confirmed stronger ALKBH2 staining in LUAD than in matched normal samples (Fig. 6B). A similar expression pattern was observed in NSCLC cell lines, with higher ALKBH2 and lower ALKBH3 levels compared with BEAS-2B bronchial epithelial cells (Fig. 6C). In survival analyses, elevated ALKBH2 was associated with worse overall survival, whereas higher ALKBH3 correlated with improved prognosis (Fig. 6, D and E).

**Figure 6.**
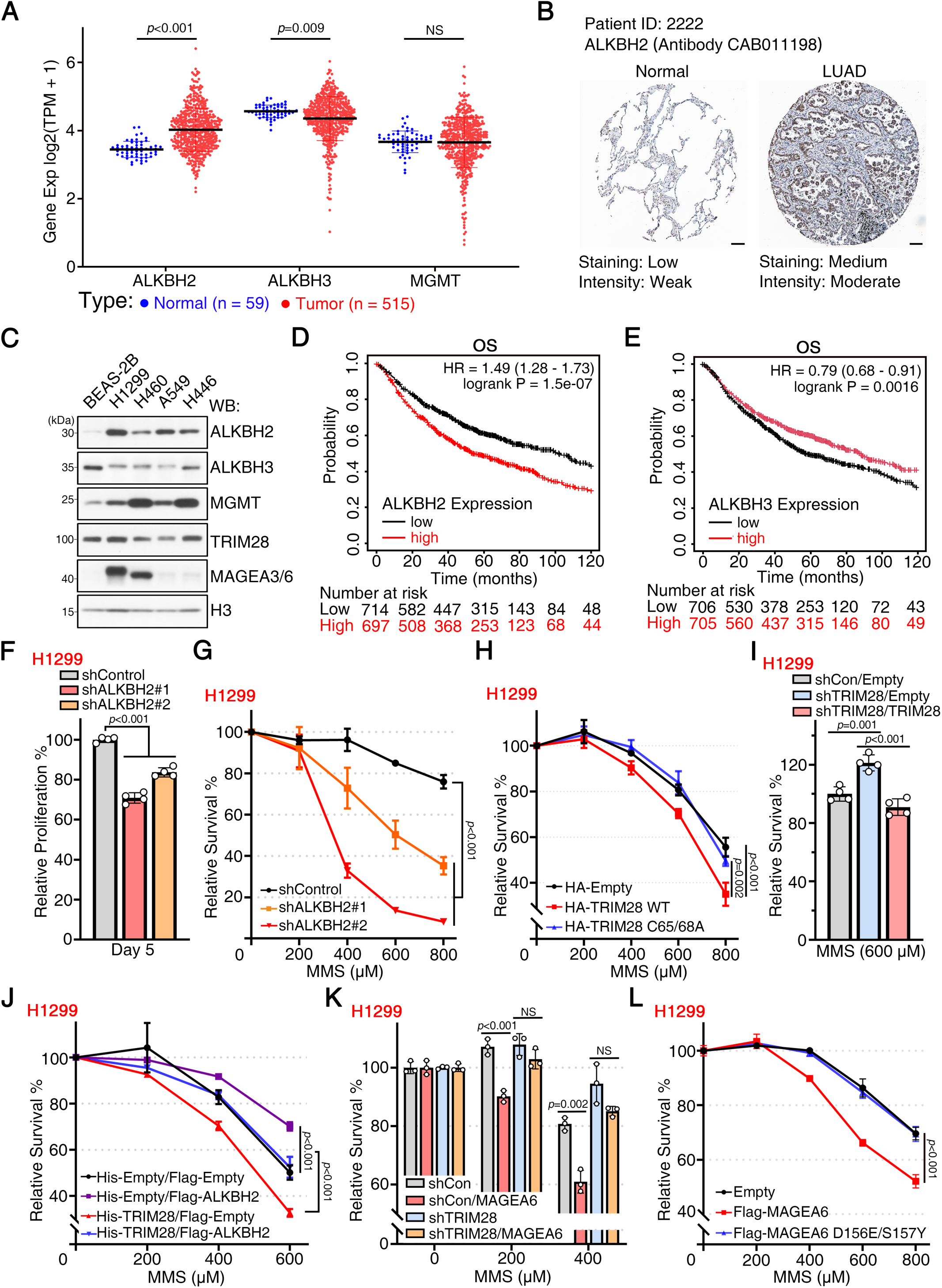
The TRIM28–ALKBH2 axis is clinically relevant and regulates sensitivity to alkylation damage. **(A)** Relative mRNA expression of ALKBH2, ALKBH3, and MGMT in LUAD versus normal lung (TCGA dataset; Mann–Whitney test). **(B)** Representative ALKBH2 IHC staining in LUAD and matched normal lung (Human Protein Atlas). Scale bar = 100 µm. **(C)** WB analysis of ALKBH2, TRIM28, ALKBH3, and MGMT in NSCLC cell lines and BEAS-2B cells. N = 3 independent experiments. **(D, E)** Kaplan–Meier overall survival of LUAD patients stratified by ALKBH2 (D) or ALKBH3 (E) mRNA expression. **(F)** CCK-8 proliferation assay in H1299 cells following ALKBH2 knockdown. **(G)** CCK-8 viability assay in H1299 cells with ALKBH2 knockdown, measured 48 h after a 1 h MMS pulse. **(H)** CCK-8 viability assay in H1299 cells overexpressing Flag-TRIM28 WT or the RING-inactive mutant (C65/68A) after MMS pulse. **(I)** MMS sensitivity assessment in H1299 cells with TRIM28 knockdown and rescue. **(J)** Rescue of MMS hypersensitivity in H1299 cells by ALKBH2 co-expression. **(K)** Effect of MAGEA6 overexpression on MMS cytotoxicity. **(L)** Effect of the TRIM28-binding–deficient MAGEA6 mutant (D156E/S157Y) on MMS sensitivity. Data points are shown for panels F–L. For panels F, G, H, I, J, and L, data points represent four technical replicates from one representative experiment; the experiment was repeated twice with similar results. Panel K contains three technical replicates and was also repeated twice with similar results. Error bars indicate mean ± SD. Statistical comparisons were performed using unpaired two-tailed Student’s *t*-tests or Mann–Whitney tests where appropriate. NS, not significant.

Because cigarette smoking generates alkylating and methylating DNA lesions (*59*), we next analyzed ALKBH2 expression according to smoking status. ALKBH2 transcript levels were higher in LUAD patients with a smoking history than in non-smokers (fig. S6B). Somatic mutations in MAGEA6, a component of the TRIM28–ALKBH2 pathway, were also more frequent in smokers, whereas MAGEA3 mutations showed no enrichment (fig. S6C). These observations suggest an association between ALKBH2 expression and smoking-linked alkylation stress.

To examine the functional relevance of these clinical associations under alkylation stress, we used an acute MMS exposure model. Because prolonged MMS treatment reduces TRIM28 and induces delayed ALKBH2 accumulation, we used a brief high-dose MMS pulse (500 μM, 1 h) that induced γ-H2AX with minimal effects on TRIM28 and ALKBH2 protein levels (fig. S6D). Under these conditions, ALKBH2 knockdown reduced cell viability and enhanced MMS sensitivity (Fig. 6, F and G). TRIM28 overexpression did not alter basal proliferation (Fig. S6E) but enhanced MMS sensitivity in a RING-dependent manner, and co-expression of ALKBH2 reversed this sensitization (Fig. 6, H-J), indicating that ALKBH2 acts downstream of TRIM28 during acute alkylation stress.

We next examined whether MAGEA6 modulates cellular sensitivity to alkylation stress. MAGEA6 overexpression increased MMS sensitivity, whereas its knockdown reduced MMS sensitivity. Re-expression of MAGEA6 restored MMS sensitivity (fig. S6F). MAGEA6 failed to enhance MMS cytotoxicity when TRIM28 was depleted, and a TRIM28-binding–deficient mutant (D156E/S157Y) exhibited a reduced sensitizing effect (Fig. 6, K and L). In contrast, MAGEA6 did not affect responses to cisplatin (CDDP) or the proteasome inhibitor bortezomib (fig. S6, G and H), indicating that its regulatory effect is preferentially associated with alkylation stress rather than broad cytotoxic stress. Collectively, these data indicate that MAGEA6 functions within the TRIM28–ALKBH2 axis to regulate ALKBH2 during alkylation stress.

### TRIM28–ALKBH2 axis regulates MMS–CDDP combinatorial sensitivity

To evaluate the clinical relevance of TRIM28, we analyzed 771 NSCLC cases. Among patients who received systemic therapy, platinum-based regimens, predominantly cisplatin-containing combinations, represented the major treatment modality (fig. S7A). High TRIM28 expression was associated with worse overall survival in chemotherapy-treated patients, whereas no prognostic association was observed in patients who did not receive chemotherapy (fig. S7, B and C). These data indicate that TRIM28 expression associates with adverse prognosis, specifically in chemotherapy-treated NSCLC patients.

To test whether TRIM28 influences cellular responses to sustained alkylation stress, we established a chronic MMS exposure model. MMS IC₅₀ values were approximately 141.5 μM in H1299 cells and 236.8 μM in H460 cells (fig. S7D). In dose–response assays, H1299 cells were exposed to MMS concentrations spanning the IC₅₀ range (40–120 μM) for 48 h, and TRIM28 depletion increased MMS resistance (Fig. 7A). In long-term clonogenic assays, H460 cells were treated with 100 μM MMS for 48 h and allowed to recover for 7 days in drug-free medium, and TRIM28 overexpression increased MMS sensitivity and reduced colony formation (Fig. 7, B and C). These results indicate that TRIM28 promotes cellular sensitivity to sustained MMS-induced alkylation stress.

**Figure 7.**
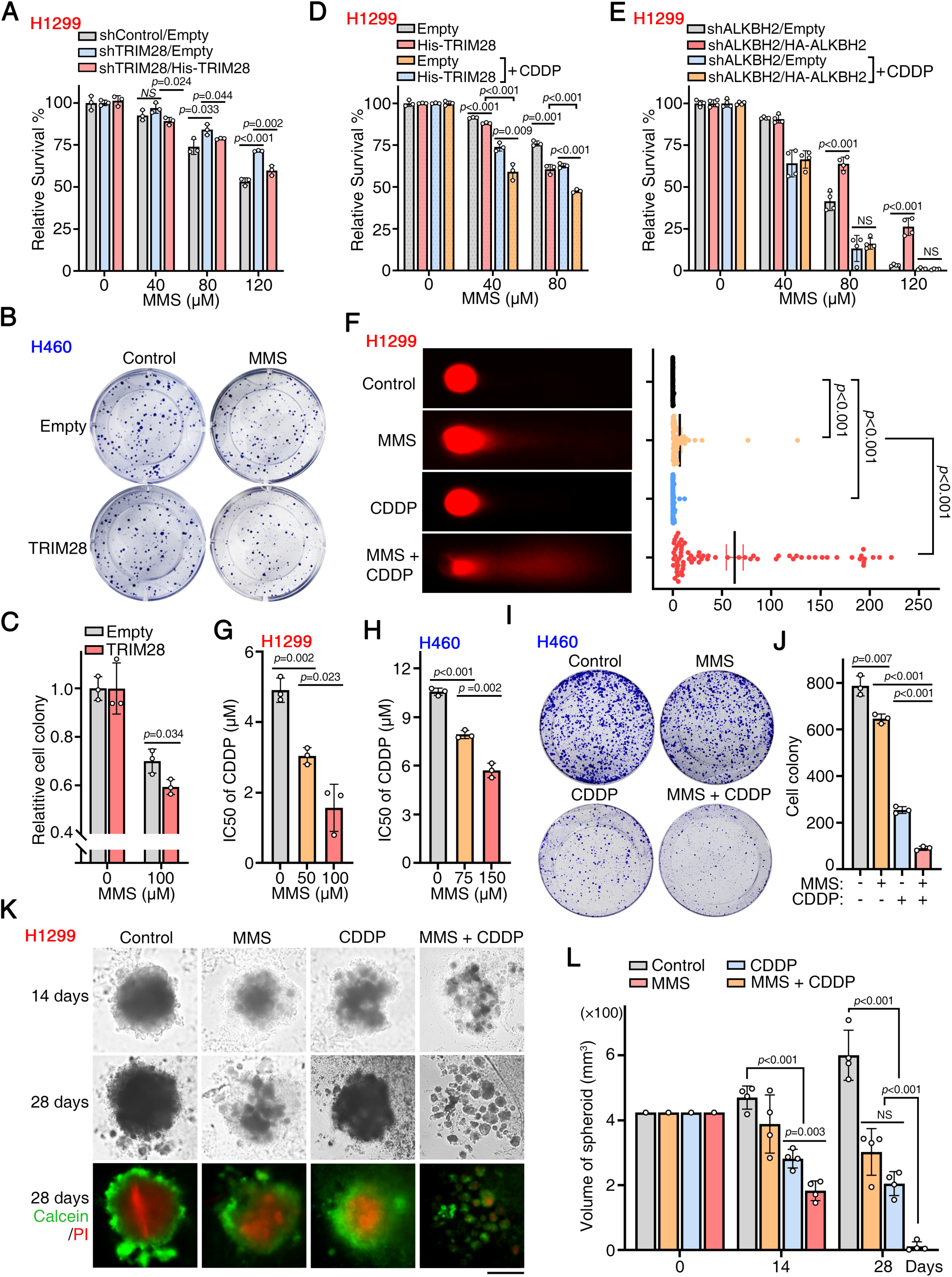
Combining MMS with CDDP synergistically enhances cancer cell killing. **(A)** CCK-8 viability assay of H1299 cells with TRIM28 knockdown treated with continuous low-dose MMS (48 h). **(B, C)** Colony formation assay of H460 cells overexpressing TRIM28 treated with MMS (100 µM) for 48 h, followed by drug removal and a 7-day growth period; representative images (B) and quantification (C). Colony numbers were averaged from 3 plates per experiment (N = 3), and the assay was repeated twice with similar results. **(D)** CCK-8 assay of H1299 cells overexpressing TRIM28 treated with MMS ± 2 µM CDDP. **(E)** CCK-8 assay of H1299 cells with ALKBH2 knockdown rescued by Flag-ALKBH2 and treated with MMS ± 2 µM CDDP. **(F)** Alkaline comet assay of H1299 cells treated with MMS (50 µM) and/or CDDP (2 µM); tail moments were quantified from 70 cells. The experiment was repeated twice with similar results. **(G, H)** IC₅₀ analysis of CDDP in H1299 (F) and H460 (G) cells with MMS co-treatment. **(I)** Representative images of clonogenic survival of H460 cells treated with MMS (75 µM) and CDDP (2 µM), and **(J)** colony quantification. Colony numbers were averaged from 3 plates per experiment (N = 3), and the assay was repeated twice with similar results. **(K, L)** 3D tumor spheroid growth of H1299 cells treated with MMS (100 µM) and CDDP (2 µM); Calcein-AM (live cells)/PI (dead cells) staining (K) and spheroid volume quantification (L). Data points represent individual spheroids from one representative experiment (n = 4 per condition); spheroid assays were independently repeated twice with similar results. Scale bar = 200 µm. For CCK-8 and IC₅₀ assays (A, D, E, G, H), data points represent technical replicates from one representative experiment (three wells per condition, four wells in panel E); each experiment was independently repeated twice with similar results. Error bars indicate mean ± SD. Statistical comparisons were performed using unpaired two-tailed Student’s *t*-tests; comet assay data in (E) were analyzed using the Mann–Whitney test. NS, not significant.

We next examined whether the TRIM28–ALKBH2 axis modulates cellular responses to combined alkylating and platinum stress. For MMS–CDDP co-treatment, 2 μM CDDP was used as a low-background dose because it was minimally cytotoxic as a single agent. TRIM28 overexpression reduced cell viability under MMS treatment, and MMS–CDDP co-treatment further decreased viability (Fig. 7D). To assess the contribution of ALKBH2, we examined MMS and CDDP sensitivity in cells with stable ALKBH2 knockdown to avoid confounding effects of MMS-induced ALKBH2 upregulation. Re-expression of ALKBH2 restored MMS tolerance, consistent with its role in alkylation repair. However, under MMS–CDDP co-treatment, low-dose CDDP attenuated ALKBH2-mediated protection and increased MMS-induced cell death (Fig. 7E). These results indicate that combined alkylating and platinum stress weakens ALKBH2-dependent MMS tolerance in a TRIM28-dependent manner.

To explore mechanisms underlying reduced ALKBH2-mediated protection during MMS–CDDP co-treatment, we assessed oxidative stress and DNA damage readouts. Because cisplatin can induce reactive oxygen species, including H₂O₂ (*60*), we tested whether H₂O₂-mediated oxidative stress enhances MMS cytotoxicity. Co-treatment of MMS with increasing concentrations of H₂O₂ reduced cell viability in a dose-dependent manner (fig. S7E), consistent with the enhanced cytotoxicity observed under MMS–CDDP co-treatment. Alkaline comet assays indicated that low-dose CDDP (2 μM) or MMS (50 μM) alone induced limited DNA strand breaks, whereas combined MMS–CDDP treatment increased tail moments (Fig. 7F) and γ-H2AX accumulation (fig. S7F). These observations indicate that oxidative and alkylating stresses are associated with increased DNA damage during MMS–CDDP co-treatment, which may contribute to reduced ALKBH2-mediated protection.

Finally, we tested whether low-level alkylation stress alters cellular sensitivity to cisplatin. Sub-IC₅₀ MMS reduced the IC₅₀ of CDDP in both H1299 and H460 cells (Fig. 7, G and H). In colony formation assays, MMS–CDDP co-treatment further reduced clonogenic survival (Fig. 7, I and J). In three-dimensional tumor spheroids, MMS–CDDP co-treatment similarly reduced spheroid size and viability (Fig. 7, K and L). Collectively, these data indicate that the TRIM28–ALKBH2 axis regulates MMS–CDDP combinatorial sensitivity and links alkylation stress to cisplatin response in NSCLC.

## DISCUSSION

Building on our previous work identifying the deubiquitinase OTUD4 as a stabilizer of an AlkB-family demethylase (*2*), this study identifies a TRIM28–MAGEA6 ubiquitin pathway that promotes ALKBH2 degradation, thereby regulating alkylation repair function. TRIM28, together with the cancer-testis antigen MAGEA6, functions as an E3 ligase complex that promotes ALKBH2 K48-linked ubiquitination and proteasomal degradation (Fig. 1–3). Under basal conditions, the TRIM28–MAGEA6 complex limits ALKBH2 protein level, whereas ALKBH2 increases TRIM28 transcription, forming a reciprocal feedback loop that maintains steady-state demethylation capacity. Alkylation stress reorganizes this loop in a time-dependent manner: acute MMS exposure enhances TRIM28–ALKBH2 association and promotes ALKBH2 ubiquitination and degradation, whereas prolonged MMS exposure promotes TRIM28 degradation, leading to ALKBH2 stabilization, while independently promoting ALKBH2 transcription. (Fig. 8).

**Figure 8.**
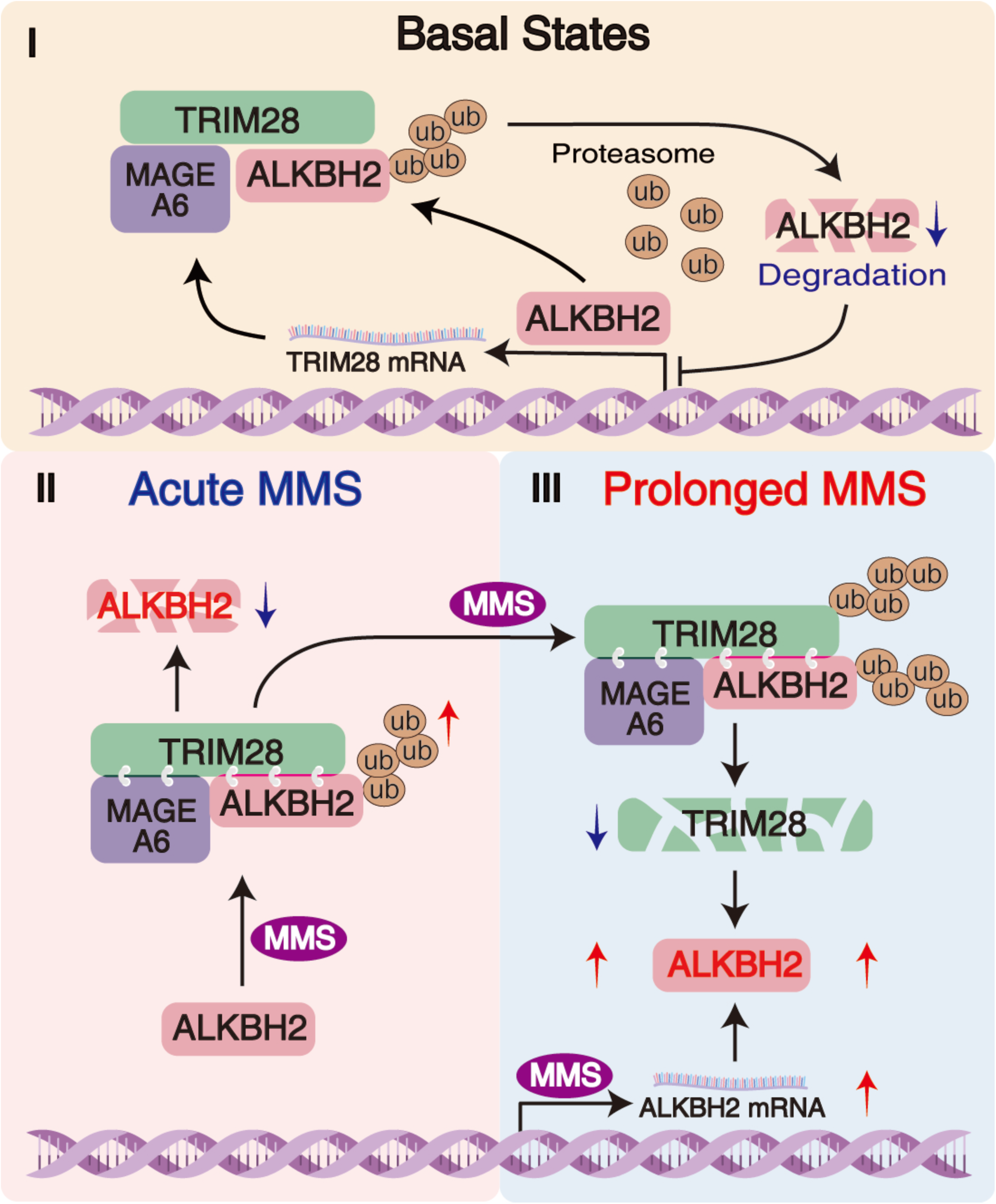
Model of the biphasic TRIM28–ALKBH2 regulatory switch induced by alkylation stress. **(I) Basal state.** MAGEA6-assisted, TRIM28-dependent K48-linked ubiquitination limits ALKBH2, while ALKBH2 increases TRIM28 transcription and protein levels. **(II) Acute alkylation stress:** MMS enhances TRIM28–ALKBH2 association, further facilitated by MAGEA6, leading to accelerated ALKBH2 ubiquitination and degradation. **(III) Prolonged alkylation stress:** Ubiquitination progressively targets TRIM28, promoting its degradation and relieving ubiquitin-dependent suppression of ALKBH2. Together with MMS-induced transcriptional upregulation of ALKBH2, this shift results in a time-dependent accumulation of ALKBH2 protein.

This study identifies the TRIM28–MAGEA6 as an E3 ligase complex that promotes K48-linked ubiquitination and degradation of ALKBH2, providing a previously uncharacterized mechanism of post-translational regulation for this duplex DNA demethylase. AlkB-family enzymes participate broadly in nucleic acid repair (*61*), and ubiquitin-dependent regulation has been characterized primarily for RNA-modifying members such as ALKBH5 and FTO (*17, 18*), whereas the pathways regulating the stability of the DNA demethylase ALKBH2, including potential E3- and DUB-linked mechanisms, remain uncharacterized. In contrast to reported MAGE–TRIM architectures (*62, 63*), MAGEA6 binds both TRIM28 and ALKBH2 (Fig. 3B), indicating a dual-binding mode that enhances complex assembly under alkylation stress. Although ALKBH2 and ALKBH3 share a conserved catalytic core, TRIM28 preferentially interacts with ALKBH2 (Fig. 1I), a selectivity likely arising from ALKBH2-specific structural features, including the finger motif and long-loop region involved in duplex DNA binding (*48*). Docking analysis maps the TRIM28-binding interface proximal to the DNA-binding region of ALKBH2 (Fig. 1J), where ssDNA competitively reduces TRIM28–ALKBH2 association (Fig. 1K), suggesting that DNA engagement transiently restricts TRIM28 access during repair and permits catalytic turnover before ubiquitin-dependent regulation. Notably, ALKBH2 increased TRIM28 mRNA and protein levels (Fig. 2, J to L), forming the transcriptional arm of this reciprocal loop, consistent with evidence that ALKBH2 modulates promoter methylation and the transcription of targets such as MGMT and BMI1 (*64, 65*). In conjunction with evidence that the deubiquitinase OTUD4 interacts with ALKBH2 and ALKBH3 (*2*), these results indicate that AlkB-family enzymes function within a broader ubiquitin-regulated network, in which the TRIM28–MAGEA6 complex contributes to demethylase homeostasis under both basal and stress conditions.

TRIM28 itself undergoes alkylation stress–dependent K48-linked ubiquitination and proteasomal degradation, introducing an additional layer of regulation within the TRIM28–ALKBH2 pathway. Notably, MMS selectively induced K48-linked polyubiquitination and proteasomal degradation of TRIM28 (Fig. 5, A to G), whereas other genotoxic agents, including TMZ, CDDP, and H₂O₂, failed to induce detectable TRIM28 degradation (Fig. 5H and fig. S5, D to F). These agents modulated ALKBH2 protein levels through TRIM28-independent mechanisms: H₂O₂ reduced ALKBH2 levels, and CDDP exerted minimal effects. In contrast, TMZ increased ALKBH2 levels via transcriptional induction, consistent with previous observations in glioblastoma cells (*10*). Mechanistically, MMS-induced TRIM28 ubiquitination and degradation were attenuated by mutation of the SUMO-site mutant K554/779/804R (Fig. 5I), suggesting that this lysine cluster contributes to TRIM28 modification under alkylation stress, potentially via SUMO-dependent regulation and/or as candidate ubiquitin-acceptor sites. Under alkylation stress, TRIM24 functioned as an upstream E3 ligase, promoting TRIM28 ubiquitination and degradation (Fig. 5, J to M, S5, H to I), thereby reversing the previously described relationship in which TRIM28 stabilizes TRIM24 (*66, 67*). This MMS-specific pathway is mechanistically distinct from RNF4-mediated TRIM28 sumoylation and degradation during double-strand break repair (*44*) and is consistent with lesion- and cue-specific organization of repair pathways and repair-complex recruitment (*68, 69*). Collectively, these results indicate that alkylation stress activates a coordinated ubiquitin-dependent regulatory program that targets both ALKBH2 and its E3 ligase, TRIM28, thereby introducing an additional layer of regulation that modulates the TRIM28–ALKBH2 axis under alkylating conditions.

A key mechanistic insight from this study is a time-dependent TRIM28–ALKBH2 feedback loop that modulates DNA repair capacity during alkylation stress. TRIM28 promotes K48-linked ubiquitination and proteasomal degradation of ALKBH2, whereas ALKBH2 increases TRIM28 expression at both mRNA and protein levels (Fig. 2, J to L), forming a regulatory loop that maintains repair homeostasis. Acute MMS exposure enhances TRIM28–ALKBH2 association and promotes ALKBH2 ubiquitination and degradation, transiently limiting repair capacity (Fig. 4). During prolonged MMS exposure, progressive TRIM28 loss is accompanied by ALKBH2 stabilization, while MMS independently induces transcriptional upregulation of ALKBH2 (Fig. 4 and 5), redirecting the loop toward higher ALKBH2 availability and repair capacity. This biphasic behavior parallels time-delayed feedback in the ATM–KAP1 (TRIM28) and p53–MDM2 circuits that coordinate DNA repair signaling with checkpoint recovery (*70–72*). Mechanistically distinct from these pathways, the TRIM28–ALKBH2 module integrates ubiquitin-mediated regulation with transcriptional feedback, enabling a homeostatic reset in which ALKBH2 restores TRIM28 levels, thereby limiting sustained ALKBH2 accumulation.

Clinically, ALKBH2 is frequently elevated in lung adenocarcinoma and associates with worse overall survival (Fig. 6, A to D), consistent with its role in removing alkylation lesions, thereby promoting tumor cell survival under genotoxic stress (*24, 73*). Notably, TRIM28 expression is associated with overall survival only in chemotherapy-treated patients, for whom higher TRIM28 levels are associated with worse overall survival (fig. S7, A to C), consistent with a treatment-dependent prognostic association. Somatic MAGEA6 variations are enriched among smokers (fig. S6C) and may further modulate this regulatory loop by altering the E3 activity of the TRIM28–MAGEA6 ubiquitin module. Given that tobacco exposure increases the endogenous burden of alkylation lesions (*74*), MAGEA6 variations may attenuate the activity of the TRIM28–MAGEA6 module and contribute to interpatient variability in alkylation repair capacity and chemoresistance. Functionally, TRIM28 sensitizes cells to MMS by promoting ALKBH2 degradation, and ALKBH2 re-expression reverses this effect (Fig. 6, F to I), indicating that ALKBH2 mediates the MMS-sensitizing effect of TRIM28. MMS–CDDP co-treatment increased DNA damage and reduced clonogenic survival, consistent with weakened ALKBH2-dependent alkylation tolerance (Fig. 7, D to K). In ALKBH2-overexpressing cells, CDDP substantially reduced ALKBH2-mediated protection, resulting in MMS sensitivity comparable to that observed upon ALKBH2 knockdown (Fig. 7E). Although TRIM28 has been linked to NSCLC chemoresistance through *miR*-125b-5p/CREB1 or E2F1 signaling (*75–79*), the present data position TRIM28 upstream of ALKBH2 degradation under alkylation stress. This context dependence is consistent with differences in lesion chemistry and stress signaling between alkylating agents and platinum crosslinkers, suggesting that the type and duration of genotoxic stress can differentially regulate TRIM28 activity. These results support a model (Fig. 8) in which a time-dependent TRIM28–ALKBH2 loop links alkylation stress to ubiquitin signaling and ALKBH2 availability, providing a conceptual framework for combination strategies in which alkylation-associated ALKBH2 induction promotes tolerance that can be counteracted by agents inducing distinct DNA lesions, such as platinum-based crosslinks.

In summary, this study identifies a stress-dependent TRIM28–ALKBH2 feedback loop that regulates alkylation repair in NSCLC cells. The TRIM28–MAGEA6 E3 complex promotes K48-linked polyubiquitination and proteasomal degradation of ALKBH2, while ALKBH2, in turn, enhances TRIM28 expression at the mRNA and protein levels, thereby maintaining repair homeostasis. Under alkylation stress, this feedback loop undergoes temporal reorganization, shifting from acute TRIM28-dependent ALKBH2 degradation to prolonged TRIM28 loss, accompanied by ALKBH2 stabilization and transcriptional upregulation. This stress-responsive feedback loop links ubiquitin-mediated control with transcription-associated regulation to coordinate cellular responses to alkylating agents. The upstream cues that trigger TRIM28 degradation, and the mechanisms by which they interface with transcriptional programs, remain to be defined. Future studies in patient-derived and *in vivo* models are needed to clarify these regulatory mechanisms and determine their relevance to chemoresistance in NSCLC.

## Methods

### Cell culture

HEK293T cells (ATCC CRL-11268, USA) and the non–small cell lung cancer (NSCLC) cell lines H1299 (SCSP-589, Cell Bank of the Chinese Academy of Sciences, China), A549 (SCSP-503, Cell Bank of the Chinese Academy of Sciences, China), and NCI-H460 (SCSP-584, Cell Bank of the Chinese Academy of Sciences, China) were maintained in Dulbecco’s modified Eagle’s medium (DMEM; C11995500BT, Gibco, USA) supplemented with 10% heat-inactivated fetal bovine serum (FBS; 10099-141C, Gibco, USA) and penicillin–streptomycin. Cells were cultured at 37 °C in a humidified 5% CO₂ incubator, and all experiments were performed using low-passage cells (passages 5–15). All cell lines were routinely confirmed to be mycoplasma-free using a luminescence-based detection assay (C0297, Beyotime, China).

### Plasmid constructs, mutagenesis, and viral infection

Full-length human TRIM28 and ALKBH2 cDNAs were used as templates for PCR-based site-directed mutagenesis. TRIM28 deletion and point mutants, including the BCC fragment (residues 141–400) and the RING-inactive C65/68A mutant, were generated by PCR, cloned into pENTR-4, and transferred into pHAGE-CMV-Flag/HA/His via Gateway™ LR recombination (11791020, Invitrogen, USA). ALKBH2 variants were cloned into pCMV-Flag for mammalian expression and into pET28a-His-2×Flag for bacterial expression. All constructs were verified by Sanger sequencing. Wild-type MAGEA6 cDNA was generated by reverse transcription of total RNA from A549 cells, and H1299-derived MAGEA6 variants were generated analogously. Transient transfections were performed using Lipo8000 (C0533, Beyotime, China).

Lentiviruses were produced in HEK293T cells by co-transfecting pLKO.1 or pHAGE vectors with the packaging plasmids psPAX2 and pMD2.G. Viral supernatants were collected 48–72 h post-transfection, clarified, and passed through 0.45-µm filters. Target cells were infected in the presence of polybrene (6 µg/mL) for 12 h. Stable cell lines were selected with puromycin (1 µg/mL) or blasticidin (5 µg/mL) for 3 days and maintained under selection-free conditions unless otherwise indicated.

### Immunoprecipitation and western blotting

For overexpression immunoprecipitation (IP), HEK293T cells were transfected with the indicated plasmids using Lipo8000 and cultured for 48–72 h before lysis. Cells were lysed in a buffer containing 150 mM NaCl, 50 mM Tris-HCl (pH 7.9), 1% Triton X-100, 10% glycerol, and 2 mM DTT, supplemented with protease and phosphatase inhibitors. Clarified lysates were incubated overnight at 4 °C with anti-Flag M2-agarose beads (A2220, Sigma-Aldrich, USA) or anti-HA-agarose beads (B26201, Selleck, China). For denaturing IP, lysates were boiled in TBS containing 1% SDS and subsequently diluted with lysis buffer to a final SDS concentration of 0.1% before IP. Each co-IP experiment was performed at least twice independently, and IP samples were analyzed by western blotting (WB) two to three times with consistent results.

For endogenous IP in H1299 cells, lysates were prepared using the same buffer and cleared by centrifugation (15,000 × *g*, 10 min, 4 °C). Supernatants were pre-cleared with Protein A/G magnetic beads (B23202, Selleck, China) for 1 h and incubated overnight at 4 °C with anti-TRIM28 antibody (1.5 µg). Immune complexes were captured with Protein A/G magnetic beads, washed with lysis buffer, and eluted in Laemmli sample buffer. Input and flow-through fractions were analyzed in parallel by WB to verify expression and loading.

For ssDNA competition assays, purified Flag-ALKBH2 was added to HEK293T whole-cell lysates supplemented with synthetic single-stranded DNA (ssDNA) oligonucleotides and incubated overnight at 4 °C. Disruption of the TRIM28–ALKBH2 interaction was assessed by Flag IP followed by WB.

For WB, proteins were separated by SDS–PAGE and transferred to PVDF membranes. Membranes were blocked with 5% non-fat milk in TBST for 1 h at room temperature, incubated with primary antibodies overnight at 4 °C, and then with HRP-conjugated secondary antibodies for 1–1.5 h at room temperature. Signals were detected using enhanced chemiluminescence (ECL). Information on antibodies used in this study is provided in table S5.

### shRNA and CRISPR/Cas9 knockdown/knockout

For shRNA-mediated knockdown, annealed oligonucleotides encoding target-specific sequences were cloned into the pLKO.1 vector (8453, Addgene, USA) and packaged into lentiviral particles using HEK293T cells. Target cells were transduced and selected with puromycin for 3 days to establish stable knockdown lines. Knockdown efficiency was confirmed by WB. A non-targeting shRNA was used as a control.

For CRISPR/Cas9-mediated knockout of TRIM28, single-guide RNAs (sgRNAs) were designed according to established criteria and cloned into the lentiCRISPRv2 vector (52961, Addgene, USA). Lentiviral particles were produced in HEK293T cells and used to transduce target cells, followed by puromycin selection. Single-cell clones were isolated by limiting dilution and expanded in 96-well plates, and successful knockout was verified by WB. All oligonucleotide sequences are listed in table S3.

### *In vitro* MBP pull-down assay

Recombinant MBP-tagged TRIM28 (MBP-TRIM28), MBP-MAGEA6, His-Flag-tagged ALKBH2 (His-Flag-ALKBH2), and MBP alone were expressed in *E. coli* Rosetta (DE3). Protein expression was induced with IPTG, and bacterial pellets were collected, lysed, and purified on amylose resin.

For pull-down assays, purified MBP-TRIM28, MBP-MAGEA6, or MBP alone was immobilized on amylose resin (E8021S, NEB, USA) and washed with binding buffer. The resin was incubated with purified His-Flag-ALKBH2 at 4 °C with gentle mixing. After thorough washing, bound proteins were eluted in Laemmli sample buffer at 95 °C and analyzed by WB using anti-His or anti-Flag antibodies.

### Immunofluorescence (IF)

Cells grown on glass coverslips were treated as indicated, fixed in 4% paraformaldehyde (P0099, Beyotime, China) for 30 min, and permeabilized with 0.5% Triton X-100 in PBS for 15 min at room temperature. After washing in PBS containing 0.1% Tween-20 (PBST), samples were incubated with primary antibodies overnight at 4 °C. The next day, coverslips were washed and incubated with fluorophore-conjugated secondary antibodies together with Hoechst 33342 or DAPI for 1.5 h at room temperature. Coverslips were mounted and imaged using an Olympus fluorescence microscope. Colocalization analysis was performed in ImageJ, and Pearson correlation coefficients were calculated when applicable.

### Cycloheximide chase assay

For cycloheximide (CHX) (2112, CST, USA) chase assays, cells were treated with 150 µg/mL CHX for the indicated durations. Cells were harvested at each time point and analyzed by WB to evaluate protein stability.

### RNA isolation and qRT-PCR

Total RNA was isolated using the RNAeasy™ RNA Isolation Kit (R0026, Beyotime, China). RNA concentration and purity were assessed using a NanoDrop spectrophotometer (Thermo Fisher Scientific) by measuring absorbance at 260 nm and determining the A260/A280 and A260/A230 ratios. First-strand cDNA was synthesized using the PrimeScript™ RT reagent Kit (RR047A, Takara, Japan), and qPCR was performed using the SYBR Green qPCR mix (TSE501, Tsingke, China). Gene expression was normalized to β-actin. Primer sequences are listed in table S4.

### Protein–protein docking and structural modeling

The tertiary structures of human ALKBH2 (PDB ID: 3BTZ), ALKBH3 (PDB ID: 2NCU), and the RBCC domain of TRIM28 (PDB ID: 6QAJ) were retrieved from the Protein Data Bank (http://www.rcsb.org). Protein–protein docking simulations were performed using the GRAMM-X web server (http://gramm.compbio.ku.edu). Predicted interfaces between TRIM28 and MAGEA6 were generated using AlphaFold3 (https://alphafoldserver.com/). Docking models were visualized and inspected using PyMOL (version 2.6.0a0).

### Mass spectrometry

TRIM28 complexes from H1299 cells stably expressing TAP-TRIM28 were immunoaffinity-purified using anti-Flag M2-agarose beads and eluted with Flag peptide (F3290, Sigma, USA). Eluates were separated by SDS–PAGE, visualized by silver staining, and subjected to LC–MS/MS. Mass spectrometry analysis was performed on an Easy-nLC 1200 system (Thermo Fisher Scientific) coupled to a Dr. Maisch GmbH C18 column (75 µm × 150 mm, 3 µm). Data-dependent acquisition (DDA) was carried out on a Q Exactive HF-X mass spectrometer (Thermo Fisher Scientific). Peptide spectra were searched against the UniProt Homo sapiens Reference Proteome (UP000005640; 81791-20230317.fasta).

### Affinity purification of K48-linked polyubiquitinated proteins (Halo-TUBE)

K48-linked polyubiquitinated proteins were affinity-purified under denaturing conditions using Halo-tagged tandem ubiquitin-binding entity (Halo-TUBE) beads. Briefly, cells were harvested, lysed, and denatured by boiling in TBS containing 1% SDS, then diluted with lysis buffer to reduce SDS to 0.1% before affinity purification. Extracts were incubated with HaloLink resin (G1915, Promega, USA) pre-bound to purified Halo-TUBE protein (10 μg per sample) overnight at 4 °C with rotation. The resin was washed five times with washing buffer, and bound proteins were eluted by boiling in Laemmli sample buffer and analyzed by WB.

### Cell proliferation, CDDP sensitivity, and synergy assays

For the methyl methanesulfonate (MMS; 129925, Sigma, USA) sensitivity assay, cells (2,000–5,000 cells per well in 96-well plates) were exposed to high-dose MMS (200–800 μM) for 1 h at 37 °C, followed by replacement with fresh medium, or were continuously treated with low-dose MMS (0–200 μM) for 48 h. Cell viability was measured at 48 h using the CCK-8 assay (K1018, ApexBio, USA).

For the cisplatin (CDDP) sensitivity assay, cells were treated with CDDP (0–40 μM) (HY-17394, MCE, USA) for 48 h, and viability was assessed using the CCK-8 assay. Half-maximal inhibitory concentration (IC₅₀) values were calculated using GraphPad Prism.

For the synergy assay, cells were treated with CDDP alone or in combination with low-dose MMS for 48 h, and viability was quantified using the CCK-8 assay.

### Colony formation assay

For clonogenic assays, H460 cells were seeded in 24-well plates (400 cells per well) and allowed to adhere overnight. The following day, cells were treated with DMSO (vehicle control) or with MMS at the indicated concentrations. For combination treatment, the medium was replaced after 48 h of MMS alone, CDDP alone, or MMS–CDDP co-treatment. Cells were then cultured for 5–10 days to allow colony formation, fixed in 4% paraformaldehyde for 10 min, stained with 0.1% crystal violet (C0121, Beyotime, China), washed, air-dried, and counted.

### Alkaline comet assay

Alkaline single-cell gel electrophoresis (comet) assays were performed using a Comet Assay Kit (KGA1302, Keygentec, China) according to the manufacturer’s instructions. Comet images were captured using an Olympus fluorescence microscope. Tail DNA% and tail moment were quantified in ImageJ (version 1.54K) using the Comet Assay plugin. Tail DNA% was calculated as: Tail DNA% = [Tail DNA intensity / (Tail DNA intensity + Head DNA intensity)] × 100%; tail moment was calculated as: tail moment = tail length × Tail DNA%.

### 3D tumorsphere culture

H1299 (∼2,000 cells per drop) were cultured as hanging drops for 72 h to allow spheroid formation, then transferred to ultra-low-attachment (ULA) 96-well plates (LV-ULA002-96UW, Liver Biotech, China). Tumor spheroids were treated with MMS and/or CDDP at the indicated concentrations for 48 h. After treatment, the medium was replaced with fresh culture medium, and spheroids were maintained for 3–4 weeks. Viability and cytotoxicity were assessed using a Calcein-AM/propidium iodide (PI) Live/Dead assay kit (C2015S, Beyotime, China). Live (green) and dead (red) fluorescence was imaged using an Olympus fluorescence microscope.

### Bioinformatics and clinical data analysis

Clinical information for TCGA LUAD and LUSC patients was obtained from the TCGA data portal (https://portal.gdc.cancer.gov/). Gene expression profiles from TCGA and protein expression data from CPTAC were analyzed using UALCAN (http://ualcan.path.uab.edu/). Survival analyses were performed using the Kaplan–Meier Plotter (http://kmplot.com/analysis) under default settings, and hazard ratios and log-rank p-values were reported. ALKBH2 immunohistochemistry (IHC) images were obtained from the Human Protein Atlas (http://www.proteinatlas.org). MAGEA6 and MAGEA3 variant data were examined using COSMIC (http://cancer.sanger.ac.uk/cosmic) and cBioPortal (http://www.cbioportal.org). Cutoffs and covariates used in each analysis are provided in the corresponding figure legends.

### Statistics and reproducibility

All statistical analyses were performed using GraphPad Prism (version 9.5.0). Data are presented as mean ± standard deviation (SD). All experiments were independently repeated at least two or three times with similar results, as specified in the figure legends. Two-group comparisons were performed using two-tailed unpaired Student’s t-tests, unless otherwise indicated, and Mann–Whitney tests were applied to nonparametric datasets. Log-rank tests were used for survival analyses, and Pearson correlation coefficients (PCCs) were calculated for colocalization analyses. IC₅₀ values were derived using four-parameter logistic regression. Fisher’s exact test was used to assess differences in MAGEA3/6 variant frequencies. Exact sample sizes, numbers of independent experiments, and statistical tests are provided in the figure legends.

## Data availability

The data supporting the findings of this study are available within the Article and its Supplementary Information.

## Acknowledgments

This work was supported by the National Natural Science Foundation of China (grant no. 31971232) and the Natural Science Foundation of Hunan Province (grant no. 2019JJ40185) to YZ.

## Author contributions

Conceptualization: YZ. Methodology: ZS, YD, YZ. Investigation: ZS, YD, YL, LHW, LYZ, XRF, LM, YS, BC, YH, YL. Data curation: ZS, YD, LYZ, XRF, LM. Formal analysis: YD, ZHL, JBL. Visualization: ZS, ZHL, JBL, YZ. Writing—original draft: ZS. Writing—review & editing: YZ. Supervision: YZ. Funding acquisition: YZ.

